## Supplementary material for "Defining human mesenchymal and epithelial heterogeneity in response to oral inflammatory disease": Text Information

### SUPPLEMENTARY INFORMATION

SUPPLEMENTARY TABLES

SUPPLEMENTARY TABLE 1. Cluster markers (all, stromal, epithelial)

SUPPLEMENTARY TABLE 2. Gene Set Enrichment Analyses (Stromal)

SUPPLEMENTARY TABLE 3. Gene Set Enrichment Analyses (Epithelial)

SUPPLEMENTARY FIGURES

SUPPLEMENTARY FIGURE 1. Single-cell profiling of healthy human gingiva datasets using 10x Chromium, Related to Figure 1.

SUPPLEMENTARY FIGURE 2. Single-cell profiling of healthy and disease human gingiva using 10x Chromium, Related to Figure 1.

SUPPLEMENTARY FIGURE 3. Re-clustering of human stromal gingival cells in health and disease, Related to Figure 3.

SUPPLEMENTARY FIGURE 4. Re-clustering of human epithelial gingival cells in health and disease, Related to Figure 4.

SUPPLEMENTARY FIGURE 5. Flow Cytometry Gating Strategies on Human Gingival Cells.

**Supplementary Figure 1. Single-cell profiling of healthy human gingiva datasets using 10x Chromium, Related to Figure 1.**

A. UMAP visualisation of human gingiva clusters from healthy human donors.
B. Scatter plots showing differential expressed genes across the two healthy samples. Panels A-B, n=2 individuals.

Figure S1

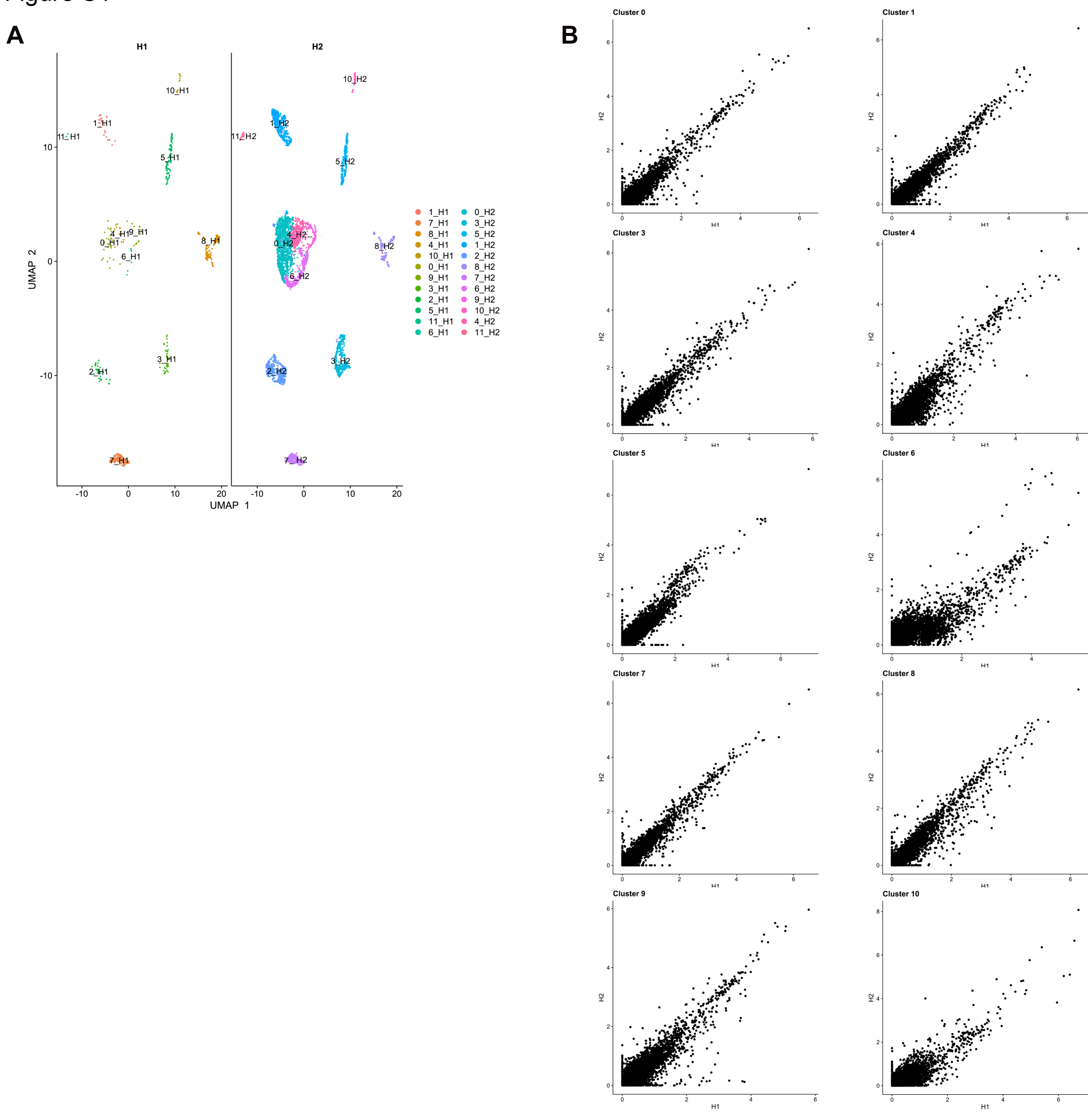

**Supplementary Figure 2. Single-cell profiling of healthy and disease human gingiva using 10x Chromium, Related to Figure 1.**

**A.** UMAP illustration of scRNA-seq data obtained from healthy and periodontitis cells (n= 12,411) from four donors coloured by condition.
**B.** Feature Plot showing the expression of lineage marker genes used for cell-type classification.

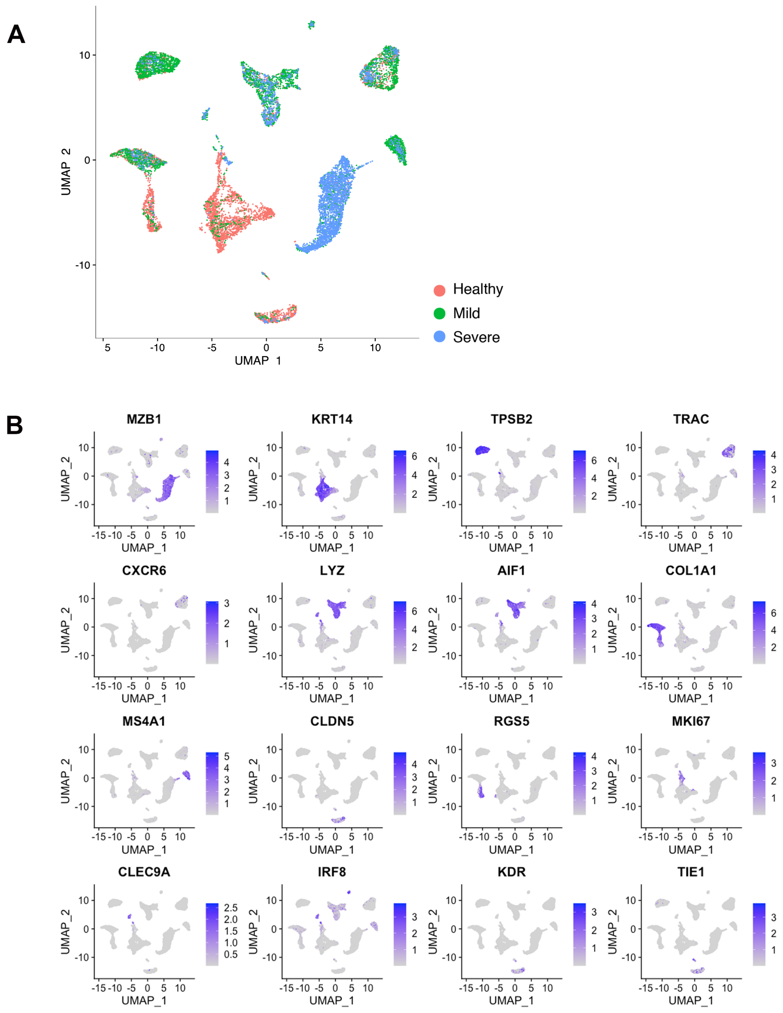

**Supplementary Figure 3. Re-clustering of human stromal gingival cells in health and disease, Related to Figure 2.**

**A.** Feature Plots showing the expression of individual genes used for cell-type assignment of different stromal subsets.

**B.** GO enrichment terms for the different stromal subsets. -log adjusted p-value shown (dotted line corresponds to FDR = 0.05).

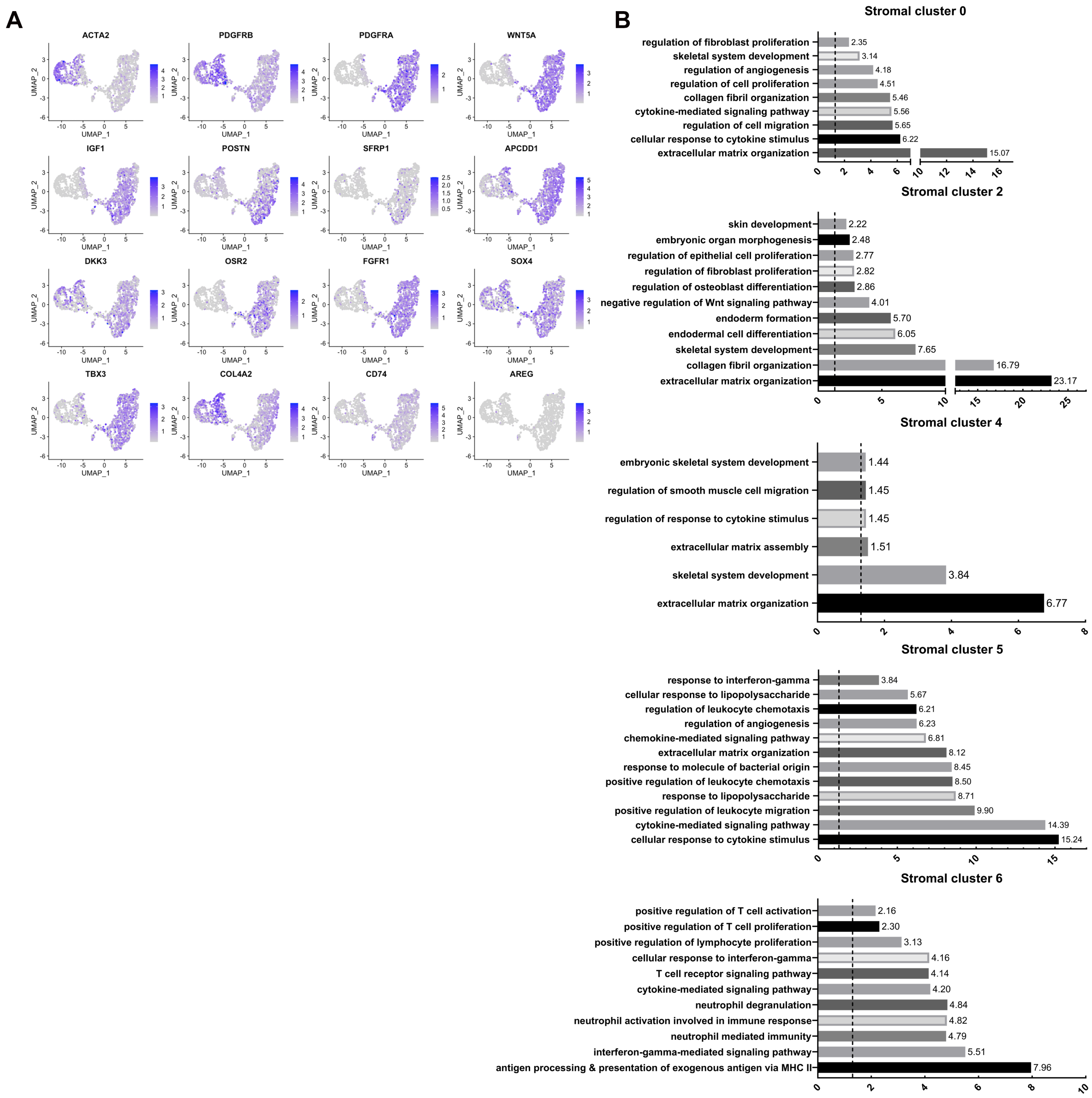

**Supplementary Figure 4. Re-clustering of human epithelial gingival cells in health and disease, Related to Figure 3.**

**A.** Feature Plots showing the expression of individual genes used for cell-type assignment of different epithelial subsets.

**B.** GO enrichment terms for the different epithelial subsets. -log adjusted p-value shown (dotted line corresponds to FDR = 0.05).

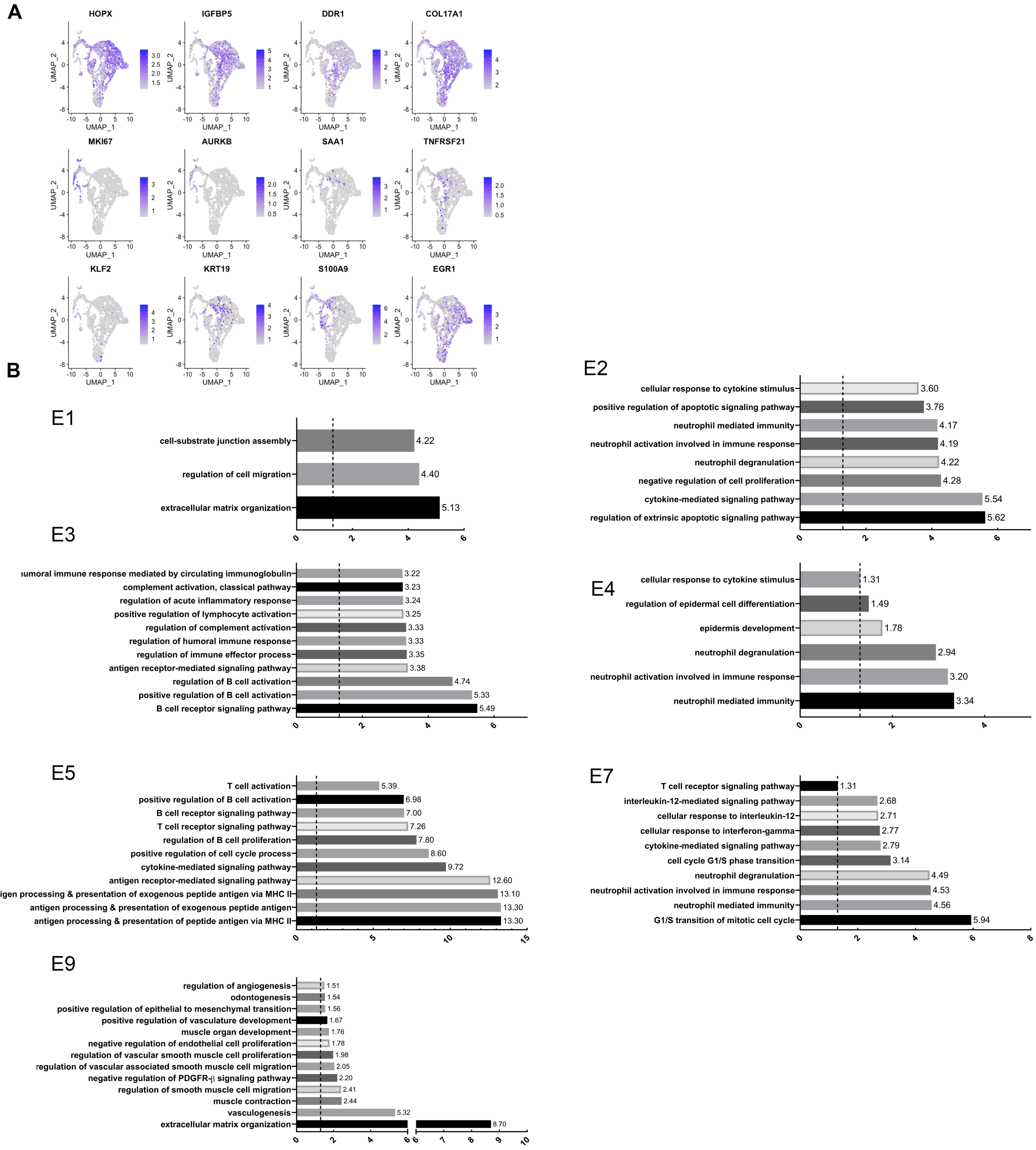

**Supplementary Figure 5. Flow Cytometry Gating Strategies on Human Gingival Cells.**

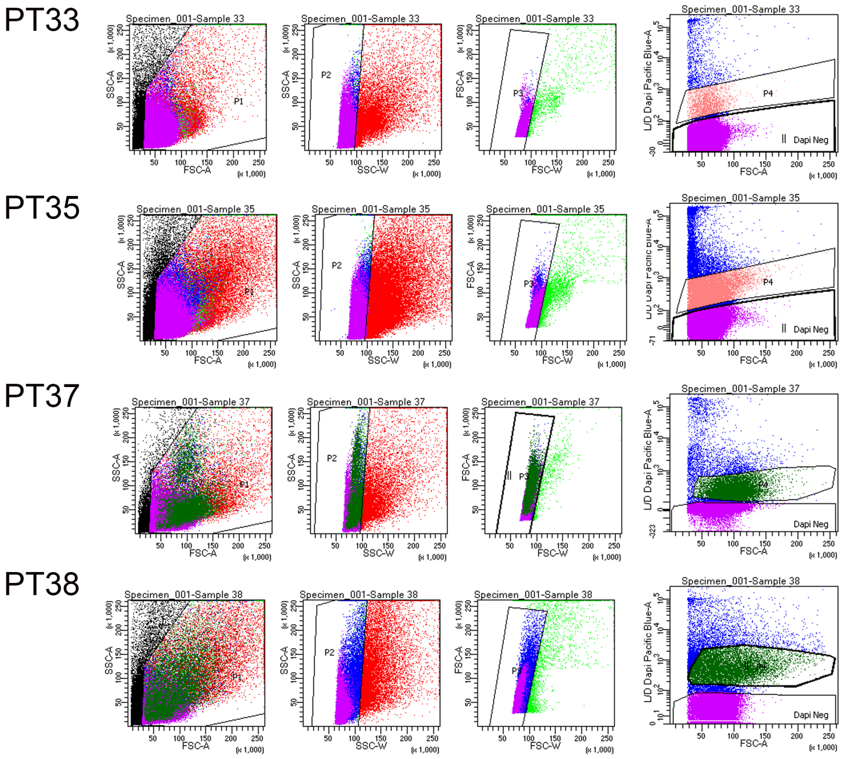
